# A new calcium-activated dynein adaptor protein, CRACR2a, regulates clathrin-independent endocytic traffic in T cells

**DOI:** 10.1101/354498

**Authors:** Yuxiao Wang, Walter Huynh, Taylor D. Skokan, Ronald D. Vale

## Abstract

Cytoplasmic dynein is a microtubule minus-end-directed motor that transports numerous intracellular cargoes. Mammalian dynein transport is initiated by coiled-coil adaptor proteins that 1) join dynein and its co-factor dynactin into a complex capable of processive motility, and 2) interact with a cargo-bound receptor, which is frequently a Rab GTPase on an organelle. Here, we report two novel dynein adaptors, CRACR2a and Rab45, which have a coiled-coil adaptor domain, a pair of EF hands, and a Rab GTPase domain fused into a single polypeptide. We find that CRACR2a-mediated dynein-dynactin motility is activated by calcium in vitro and in cells. In activated T cells, CRACR2a localizes to clathrin-independent endosomes that require microtubule-based transport to detach from the actin cortex and travel towards the microtubule organizing center. Together these results represent the first known examples of Rab GTPases that directly act as dynein adaptors and implicate CRACR2a-dynein in regulation of endocytic trafficking in T cells.

## Introduction

Microtubule-based transport is essential for the positioning of large membrane organelles, the trafficking of small transport vesicles, and the localization of mRNA and protein (Vale, 2003; Schliwa and Woehlke, 2003). Microtubules are polarized filaments with distinct plus and minus ends; in most cells, microtubule plus ends extend to the cell periphery and minus ends are anchored to the microtubule organization center (MTOC). Plus-end-directed transport is mediated by kinesin, a large family of motor proteins with over 40 members in mammals (Hirokawa et al., 2009). In contrast, all minus-end-directed intracellular transport in animal cells is carried out by a single motor protein complex, cytoplasmic dynein-1 (referred to as dynein hereafter) (Vale, 2003; Reck-Peterson et al., 2018). Most, if not all, membrane organelles are transported by dynein (Reck-Peterson et al., 2018).

The dynein holoenzyme is composed of a ~500 kD heavy chain and five smaller subunits (three light chains, one light-intermediate chain and one intermediate chain). Mammalian dynein does not display processive motility due to autoinhibition of its motor domain (Zhang et al., 2017). Binding of the dynactin complex and an adaptor protein activates dynein, enabling it to move processively along microtubules (McKenney et al., 2014; Schlager et al., 2014). Thus far, eight adaptors have been demonstrated to directly bind and activate dynein: BicDL1, BicD2, Hook1, Hook3, Rab11FIP3, Spindly, Ninein and Ninein-like protein (Reck-Peterson et al., 2018). A common feature of these proteins is the presence of long coiled-coil domains. Structural studies revealed that the coiled-coils of BicD and Hook are directly involved in joining dynein and dynactin together into a tripartite complex (Urnavicius et al., 2015, 2018).

Recruitment of dynein-dynactin-adaptor complex to specific membrane organelles is often mediated by the interaction between a dynein adaptor and a Rab GTPase, which associates with intracellular membrane compartments through C-terminal prenylation (Hutagalung and Novick, 2011; Reck-Peterson et al., 2018). Rab GTPases have been shown to regulate the localization as well as the conformation of dynein adaptors. Rab6, which localizes to Golgi-derived vesicles, recruits the dynein adaptor BicD to mediate retrograde transport (Matanis et al., 2002). Binding of Rab6 to BicD also alleviates its autoinhibition and promotes BicD mediated dynein-dynactin activation (Hoogenraad et al., 2003; Liu et al., 2013; Huynh and Vale, 2017). Thus far, no Rab has been shown to possess the ability to interact with and activate dynein directly.

Here, we report the discovery of two novel dynein adaptor proteins, Rab45 and CRACR2a, both of which are also Rab GTPases. These are the first identified dynein-dynactin adaptors that contain both a Rab GTPase domain and a coiled-coiled dynein-dynactin activator domain integrated within the same polypeptide. We find that the adaptor protein function of CRACR2a is activated by calcium elevation within a physiological range. In T cells, CRACR2a localizes to intracellular compartments that migrate towards the MTOC in response to T cell activation-induced calcium influx. We also show that CRACR2a forms distinct puncta at the plasma membrane of activated T cells, which give rise to internalized vesicles generated through clathrin-independent endocytosis. Our results demonstrate that Rab45 and CRACR2a are a new group of activating adaptors for dynein, and reveal a role of CRACR2a in dynein-mediated endocytic transport processes in T cells.

## Results

### Identification of Rab45 and CRACR2a as activating adaptors for dynein

Rab45 and CRACR2a are atypical Rab GTPases that contain a pair of EF hand domains, a coiled-coil domain, and a Rab G domain (Fig. 1A). We speculated that these proteins may function as dynein adaptor proteins due to (1) their domain architecture (EF hands followed by coiled-coil), which resembles that of known dynein adaptors, such as FIP3 and Ninein (Fig. 1A), and (2) their localization to a small, perinuclear compartment at the cell center, suggestive of microtubule minus-end-directed transport (Srikanth et al., 2016; Shintani et al., 2007). The human CRACR2a gene was shown to undergo alternative splicing to produce two isoforms (Srikanth et al., 2016; Wilson et al., 2015). In this study, we focused on the long isoform, as the short isoform lacks the Rab GTPase domain and does not seem to be involved in membrane trafficking (Srikanth et al., 2016).

**Figure 1.**
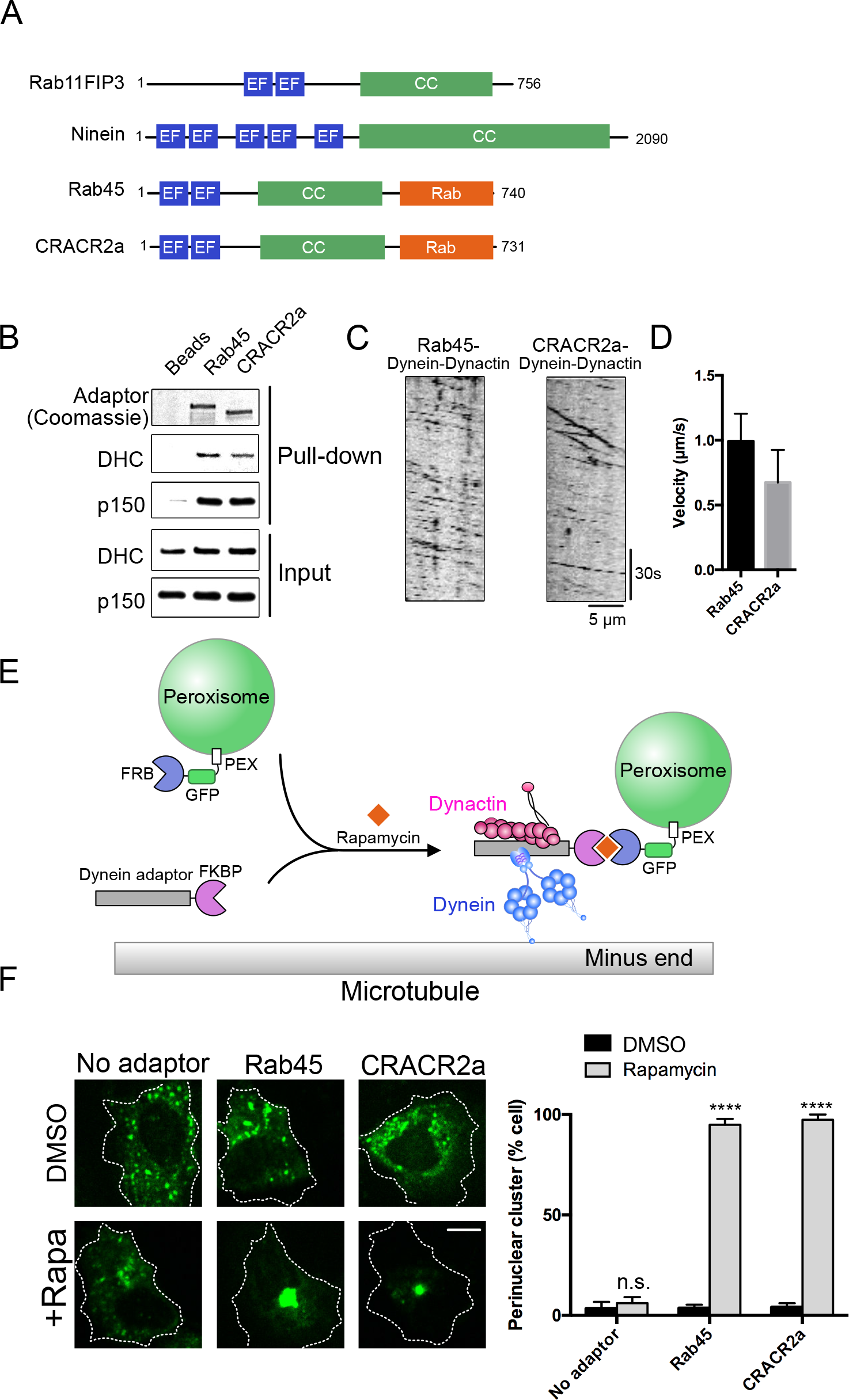
Rab45 and CRACR2a are adaptor proteins for dynein. (**A**) Domain organizations of Rab45 and CRACR2a compared to Rab11FIP3 and Ninein. EF: EF-hand; CC: coiled-coil. Sizes of the domains are not to scale. (**B**) Pull-down assay showing Rab45 and CRACR2a bind to dynein-dynactin. DHC: Dynein Heavy Chain; p150: the p150 subunit of dynactin. (**C**) Sample kymographs of microtubule-based motility of Rab45 or CRACR2a in complex with dynein-dynactin. Fluorescence is from the N-terminal GFP tag on Rab45 or CRACR2a. (**D**) Quantification of the velocities of dynein-dynactin in complex with Rab45 or CRACR2a. Error bar: SD; n = 40 processive particles from three replicates. (**E**) Schematic of the inducible peroxisome trafficking assay. U2OS cells were transfected to express GFP-FRB fused with a peroxisome targeting sequence (PEX) and an mCherry-adaptor-FKBP construct. Treatment of the cell with rapamycin induces the localization of the dynein-dynactin-adaptor complex to peroxisome and drives retrograde transport of peroxisomes towards the microtubule minus end. (**F**) Representative images of the peroxisome transport assay (scale bar: 10 *μ*m). +Rapa indicates treatment with rapamycin. Quantifications of the percentage of cells with a strong perinuclear cluster of peroxisomes are shown on the right. Error bar: SEM; n = 3 independent experiments (30-40 cells per experiment). **** p <0.0001 between DMSO-treated group and rapamycin-treated group using two-way ANOVA analysis with Sidak’s multiple comparisons test.

We first sought to determine whether Rab45 and CRACR2a could interact with dynein and dynactin. To this end, we expressed and purified full-length Rab45 and CRACR2a from bacteria (Fig. S1), attached these proteins to beads, and then performed pull-down assays with dynein-dynactin purified from human RPE1 cells. Indeed, both Rab45 and CRACR2a interacted with dynein-dynactin in these pull-down assays (Fig. 1B).

We next examined the ability of Rab45 and CRACR2a to activate the processive motility of native dynein-dynactin using an established total internal reflection fluorescence microscopy (TIRF-M) single molecule assay (McKenney et al., 2014; Huynh and Vale, 2017). Purified Rab45 and CRACR2a robustly activated the processive motility of dynein-dynactin on microtubules (Fig. 1C and 1D), similar to other known dynein adaptors (McKenney et al., 2014; Schlager et al., 2014). The velocities of movement were 0.99 ± 0.21 *μ*m/s and 0.67 ± 0.25 *μ*m/s (mean ± S.D.) for Rab45 and CRACR2a, respectively. Together, these data indicate that Rab45 and CRACR2a can bind to dynein-dynactin and activate dynein motility in vitro.

To determine whether Rab45 and CRACR2a activate dynein transport in cells, we employed a previously developed inducible peroxisome trafficking assay, in which a candidate adaptor protein is induced to localize to peroxisomes by addition of rapamycin (Kapitein et al., 2010) (Fig. 1E). In the absence of rapamycin, peroxisomes are distributed randomly throughout the cell. Recruitment of a dynein adaptor to peroxisomes causes these organelles to move towards the microtubule minus end and accumulate around the MTOC (Fig. 1E). We found that rapamycin-induced targeting of Rab45 and CRACR2a to peroxisomes resulted in strong clustering of peroxisomes at the cell center (Fig. 1F), suggesting that Rab45 and CRACR2a are capable of recruiting dynein and activating dynein-mediated transport in cells.

### Regulation of the dynein adaptor function of Rab45 and CRACR2a by calcium

The presence of EF-hands, a known calcium binding motif, in Rab45 and CRACR2a led us to examine whether calcium regulates their function as dynein adaptors. In the previous pull-down and single-molecule assays (Fig. 1B and 1C) we did not add calcium or EGTA, in which case trace amount of calcium in the 1-10 *μ*M range could still be present in the buffer (Bers et al., 2010). We therefore repeated the dynein-dynactin binding assay either with 2 mM EGTA, to deplete calcium completely, or with 2 *μ*M free calcium, a concentration relevant for most calcium-dependent physiological processes (Bers et al., 2010) (see Materials and Methods for details). While complex formation between Rab45 and dynein-dynactin was insensitive to calcium concentration, CRACR2a required the presence of calcium for stable interaction with dynein and dynactin (Fig. 2A). In addition, mutating the calcium binding sites in the EF hands of CRACR2a (CRACR2A^EFmut^) abolished its interaction with dynein-dynactin, regardless of calcium concentration (Fig. 2A).

**Figure 2.**
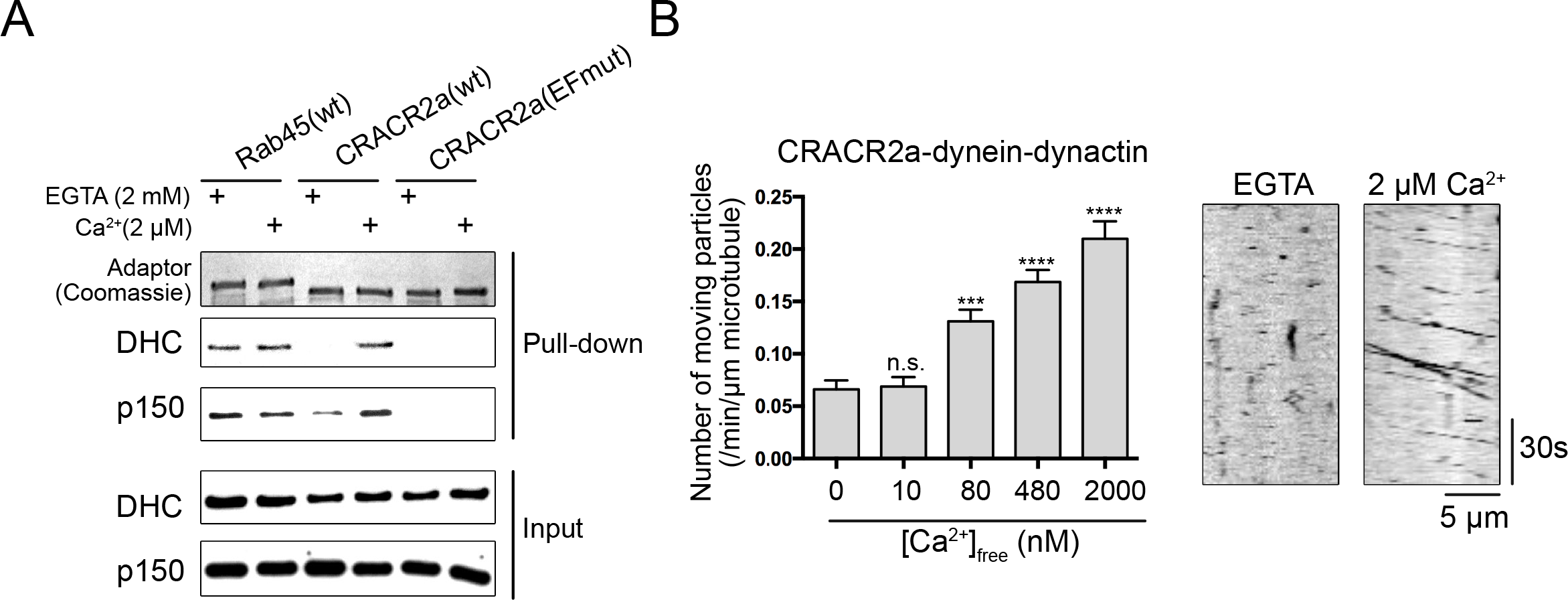
Regulation of the dynein adaptor function of CRACR2a by calcium. (**A**) Dynein-dynactin pull-down assays performed with calcium-depleted buffer (2 mM EGTA) or buffer containing 2 *μ*M free calcium. DHC: Dynein Heavy Chain; p150: the p150 subunit of dynactin. (**B**) Quantification of CRACR2a-dynein-dynactin motility at different free calcium concentrations. Data are from three replicates, each measuring at least 20 microtubules. Error bar: SEM. ***p <0.001. **** p<0.0001, one way ANOVA with Dunnett’s test versus EGTA condition. Example kymographs are shown on the right.

We further tested whether the single molecule motility of CRACR2a-dynein-dynactin responds to changes of calcium in the physiological range (100 nM to ~10 *μ*M). Raising calcium concentration from 100 nM to 2 *μ*M significantly increased the number of processive events on microtubules (Fig. 2B). In contrast, the motility of Rab45-dynein-dynactin was similar in the presence of EGTA or 2 *μ*M calcium (Fig. S2), consistent with the result from the pull-down assay (Fig. 2A). Taken together, these results suggest that the dynein adaptor function of CRACR2a is activated by physiological levels of calcium increase.

### T cell activation induced calcium elevation stimulates the transport of CRACR2a towards the MTOC

We next sought to determine whether the dynein adaptor protein function of CRACR2a is regulated by calcium in cells. A previous study reported that CRACR2a is highly expressed in primary T cells as well as in the Jurkat T cell line (Srikanth et al., 2016). We observed that stably-expressed GFP-CRACR2a in Jurkat cells localized to a loosely connected perinuclear membrane compartment as well as to small vesicles that traveled bidirectionally along microtubules (Fig. 3A, Video 1). Strikingly, treatment of the cells with ionomycin, which elevates cytoplasmic calcium, caused a rapid accumulation of GFP-CRACR2a at the MTOC (Fig. 3A, 3B). Mutating the calcium binding sites in CRACR2a EF hands abolished ionomycin-induced MTOC clustering of GFP-CRACR2a, consistent with the in vitro observation that CRACR2a^EFmut^ failed to bind to dynein-dynactin in response to calcium elevation. To test the notion that the calcium-stimulated accumulation of CRACR2a at the MTOC is caused by an increase of retrograde transport of CRACR2a vesicles, we expressed a GFP-CRACR2a mutant lacking the C-terminal polybasic region and prenylation motif (GFP-CRACR2a^Δtail^). As expected, GFP-CRACR2a^Δtail^ was diffusive in the cytoplasm (Fig. 3A). Moreover, ionomycin treatment failed to induce clustering of GFP-CRACR2a^Δtail^ at the MTOC (Fig. 3B), suggesting that membrane association is essential for the observed accumulation of CRACR2a at the MTOC in response to calcium elevation.

**Figure 3.**
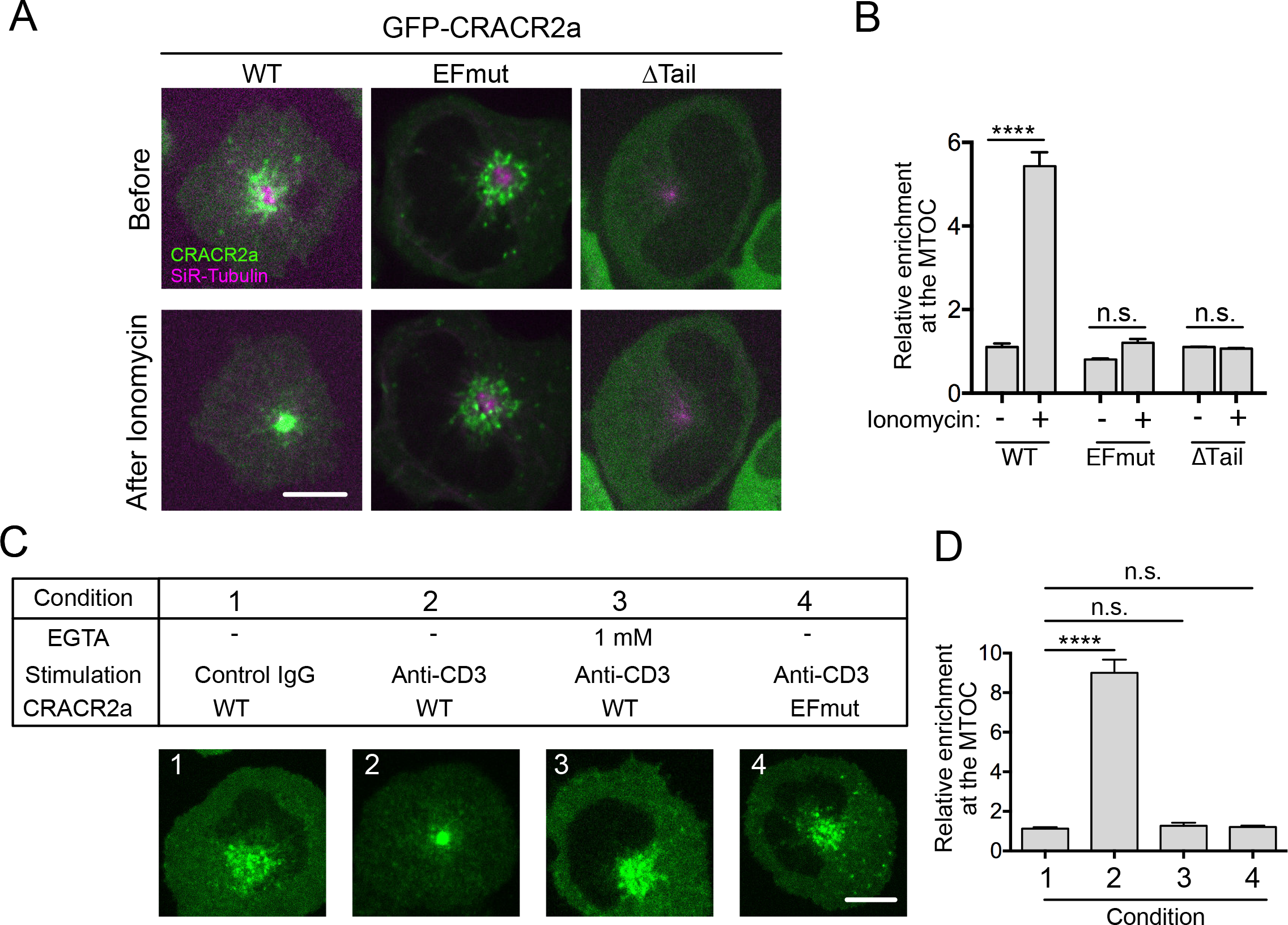
Calcium regulation of CRACR2a subcellular localization in T cells. (**A**) Subcellular distribution of CRACR2a in response to ionomycin stimulation. The same cell before and after ionomycin treatment is shown. MTOC is visualized by staining with 200 nM SiR-tubulin for 4 hr. Scale bar: 10 *μ*m. EFmut indicates mutated calcium binding site in EF hands (D63A, E65A, D97A, D99A). ΔTail indicates deletion of C-terminal prenylation tail (Δ725-731). (**B**) Quantifications of relative enrichment of CRACR2a at the MTOC. n = 20-30 cells from three independent experiments. Error bar indicates SEM. **** indicates p <0.0001, two way ANOVA with Sidak’s multiple comparisons test. (**C**) T cell activation induces clustering of CRACR2a compartments at the MTOC. Scale bar: 10 *μ*m. (**D**) Quantifications of the levels of relative enrichment at MTOC for data shown in (C). n = 20-30 cells from three independent experiments. Error bar indicates SEM. **** p <0.0001, one way ANOVA with Dunnett’s test comparing each sample with sample #1.

T cell receptor (TCR) activation induces a rapid increase in intracellular free calcium to initiate downstream signaling. To determine if CRACR2a dynamics are altered during TCR triggering, we stimulated Jurkat cells expressing GFP-CRACR2a with immobilized anti-CD3ε antibody (clone OKT3). T cell activation induced a rapid congregation of GFP-CRACR2a at the MTOC (Fig. 3C and 3D). Depletion of calcium from the media, as well as disruption of the calcium binding sites in the EF hands of CRACR2a, abolished the ability of TCR activation to cause CRACR2a clustering at the MTOC (Fig. 3C and 3D). These results indicate that CRACR2a responds to calcium influx induced by TCR activation and drives dynein-mediated transport towards the MTOC.

We sought to determine whether the CRACR2a compartment at the MTOC colocalizes with known membrane organelles. An earlier study suggested that CRACR2a colocalized with Rab8 positive trans-Golgi network in T cells (Srikanth et al., 2016). Although GFP-CRACR2a and mCherry-Rab8 in Jurkat cells both localized to perinuclear region in general, high resolution confocal imaging revealed little colocalization between the two proteins (Fig. S3A). In addition, ionomycin treatment did not change the distribution of Rab8A, whereas it strongly compacted CRACR2a-containing vesicles at the MTOC (Fig. S3A, S3B). We further tested the colocalization between CRACR2a and Rab11A, which localizes to perinuclear recycling endosomes. Upon ionomycin stimulation CRACR2a but not Rab11A congregated at the MTOC (Fig. S3C, S3D). These results together suggest that the subcellular localization of CRACR2a is distinct from those of Rab8 and Rab11.

### Formation and dynamics of CRACR2a puncta at the T cell immunological synapse

Activation of a T cell by an antigen-presenting cell leads to the formation of a specialized junction known as the immunological synapse (IS) (Dustin et al., 2010). The IS is not only the site of T cell signal transduction but also a hotspot of intracellular membrane trafficking (Griffiths et al., 2010; Onnis et al., 2016). We induced IS formation in vitro by stimulating Jurkat cells with immobilized anti-CD3, and examined the dynamics of CRACR2a at the IS using TIRF-M, which restricts the zone of illumination to regions adjacent to the cell-glass interface. We observed the MTOC-associated GFP-CRACR2a compartment at the center of the IS (Fig. 4A), consistent with previous reports that the MTOC moves to a location underneath the IS during T cell activation (Lowin-Kropf et al., 1998; Martín-Cófreces et al., 2008).

**Figure 4.**
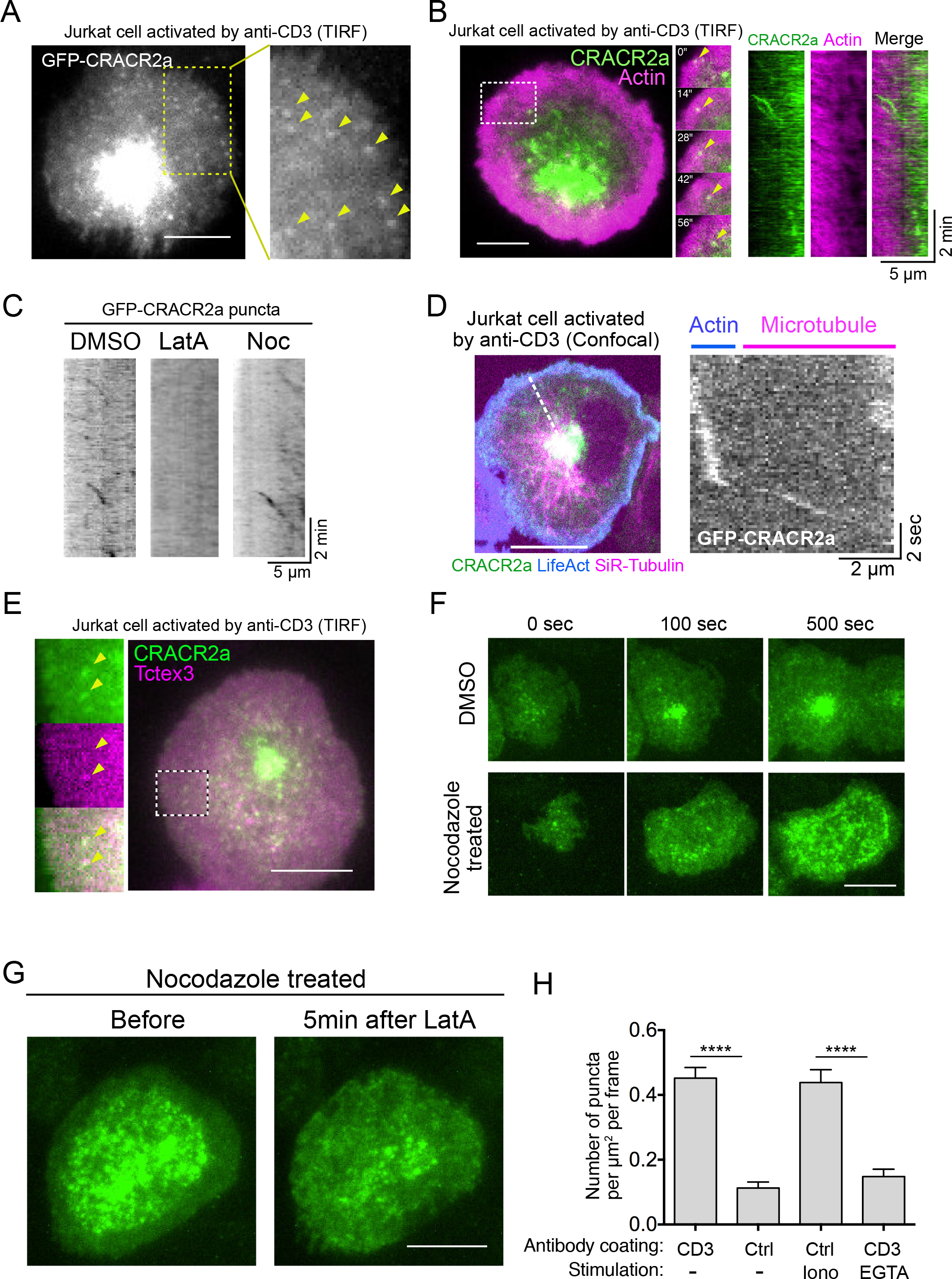
Formation and dynamics of CRACR2a puncta at the T cell immunological synapse. (**A**) TIRF-M imaging reveals formation of CRACR2a puncta (indicated by yellow arrow heads) at the periphery of the immunological synapse. Scale bar: 10 *μ*m. (**B**) CRACR2a puncta migrate with actin retrograde flow. Scale bar: 5 *μ*m. Kymograph of CRACR2a puncta and actin is shown on the right. (**C**) Sample kymographs of CRACR2a puncta dynamics at the IS periphery. Jurkat cells were treated with 2 *μ*M Latrunculin A (LatA) to depolymerize actin or 5 *μ*M Nocodazole (Noc) to depolymerize microtubules. (**D**) Imaging CRACR2a puncta dynamics on actin (visualized by BFP-LifeAct) and microtubules (visualized by SiR-Tubulin) using a spinning disk confocal microscope. Kymograph analysis along the indicated line is shown on the right. (**E**) Colocalization between CRACR2a and dynein (visualized by mCherry-tagged Tctex3, a dynein light chain). Scale bar: 10 *μ*m. (**F**) Nocodazole disrupts the transport of CRACR2a towards the MTOC and causes CRACR2a puncta to accumulate at the central regions of the IS. Scale bar: 10 *μ*m. (**G**) F-actin disassembly causes dispersal of CRACR2a puncta accumulated at the IS in nocodazole treated cells. Scale bar: 10 *μ*m. The same cell before and 5 min after Latrunculin (LatA) treatment is shown. Images are representative of 15-30 cells examined in three independent experiments. (**H**) Quantification of the amount of observed CRACR2a puncta when Jurkat cells are stimulated by different conditions. CD3 indicates anti-CD3 (OKT3); Ctrl indicates IgG isotype control. Iono indicates stimulation with ionomycin. n = 25-30 cells from three independent experiments. Error bar indicates SEM. **** p <0.0001, one way ANOVA with Tukey’s multiple comparisons test.

Intriguingly, we also observed that GFP-CRACR2a formed numerous diffraction-limited puncta that underwent centripetal movement at IS periphery (Fig. 4A, Video 2). Visualization of actin dynamics at the IS using LifeAct revealed that CRACR2a puncta co-migrated with the retrograde flow of actin (Fig. 4B, Video 3). Disruption of F-actin using Latrunculin A (2 *μ*M) rapidly abolished the movement of CRACR2a puncta (Fig. 4C, Video 4). These results suggest that actin retrograde flow drives the observed motility of CRACR2a puncta at the IS.

We noticed that CRACR2a puncta migrated centripetally for a short distance (< 5 *μ*m) and then suddenly disappeared (e.g. in Fig. 4B kymograph), indicating that they moved away from the TIRF illumination zone. We re-examined CRACR2a puncta dynamics using a spinning disk confocal microscope with fluorescent labeling of actin and microtubules, which revealed that CRACR2a puncta motility can be separated into two phases: an initial phase of slow, actin-driven motility at the cell periphery (0.06 ± 0.014 *μ*m/s, mean ± SD), followed by a later phase of fast movement on microtubules towards the MTOC (1.8 ± 0.7 *μ*m/s, mean ± SD) (Fig. 4D). Colocalization analysis confirmed that dynein was recruited to CRACR2a puncta immediately after their appearance at the distal edge of the IS (Fig. 4E). Depolymerization of microtubule by nocodazole treatment (5 *μ*M) did not affect the retrograde movement of CRACR2a puncta at the peripheral region of the IS (Fig. 4C). However, under these conditions, CRACR2a failed to form a tight cluster at the MTOC, but instead accumulated within the TIRF field (<200 nm zone near the plasma membrane) at the central region of the IS (Fig. 4F, Video 5). Further disruption of F-actin in nocodazole-treated cells caused the accumulated CRACR2a puncta to completely disperse (Fig. 4G), suggesting that CRACR2a puncta remain associated with the actin cortex when microtubule-based transport is absent. Together, the above results indicate that dynein recruitment to CRACR2a puncta play important role in their dissociation from actin cortex and initiation of microtubule-based motility.

To determine whether T cell signaling regulates the formation of CRACR2a puncta, we plated Jurkat cells onto isotype control antibody-coated glass that does not activate TCR and observed greatly reduced number of CRACR2a puncta at the cell membrane (Fig. 4H). Addition of ionomycin in this condition is sufficient to stimulate the formation of CRACR2a puncta, even in the absence of TCR proximal signaling (Fig. 4H). In contrast, depletion of extracellular calcium by EGTA, which prevents cytosolic calcium elevation upon TCR activation, blocked the formation of CRACR2a puncta when cells are activated by immobilized anti-CD3 (Fig. 4H). These results suggest that the formation of CRACR2a puncta at the IS is stimulated by TCR activation-induced calcium elevation.

### CRACR2a is involved in clathrin-independent endocytosis in T cells

The observation that CRACR2a puncta eventually travel along microtubules towards the MTOC suggests that they could mature into intracellular vesicles. This notion is consistent with a previous report that CRACR2a localizes to sub-synaptic vesicles during T cell activation (Srikanth et al., 2016). Rab5-positive early endosomes and Rab11-positive recycling endosomes have been shown to mediate vesicle trafficking at the IS (Finetti et al., 2014). We did not observe colocalization of GFP-CRACR2a puncta with either Rab5 or Rab11 (Fig. 5A). Furthermore, there was no detectable localization of GFP-CRACR2a to clathrin coated pits (Fig. 5A). These results suggest that the CRACR2a puncta are distinct from these endosomal populations. To test this further, Jurkat cells were stimulated with immobilized anti-CD3 antibodies and treated with a panel of chemical inhibitors of clathrin-coated pit formation (chlorpromazine (CPZ) or PitSTOP2), dynamin function (MiMAB) or caveolae-mediated uptake (Filipin). None of these inhibitors affected the number of observed CRACR2a puncta (Fig. 5B). In contrast, inhibition of actin polymerization (with 2 *μ*M Latrunculin A or 5 *μ*M Cytochalasin B) completely abolished the appearance of new GFP-CRACR2a puncta (Fig. 5B). These results suggest that CRACR2a vesicles form through a clathrin-independent, caveolin-independent but actin-dependent pathway.

**Figure 5.**
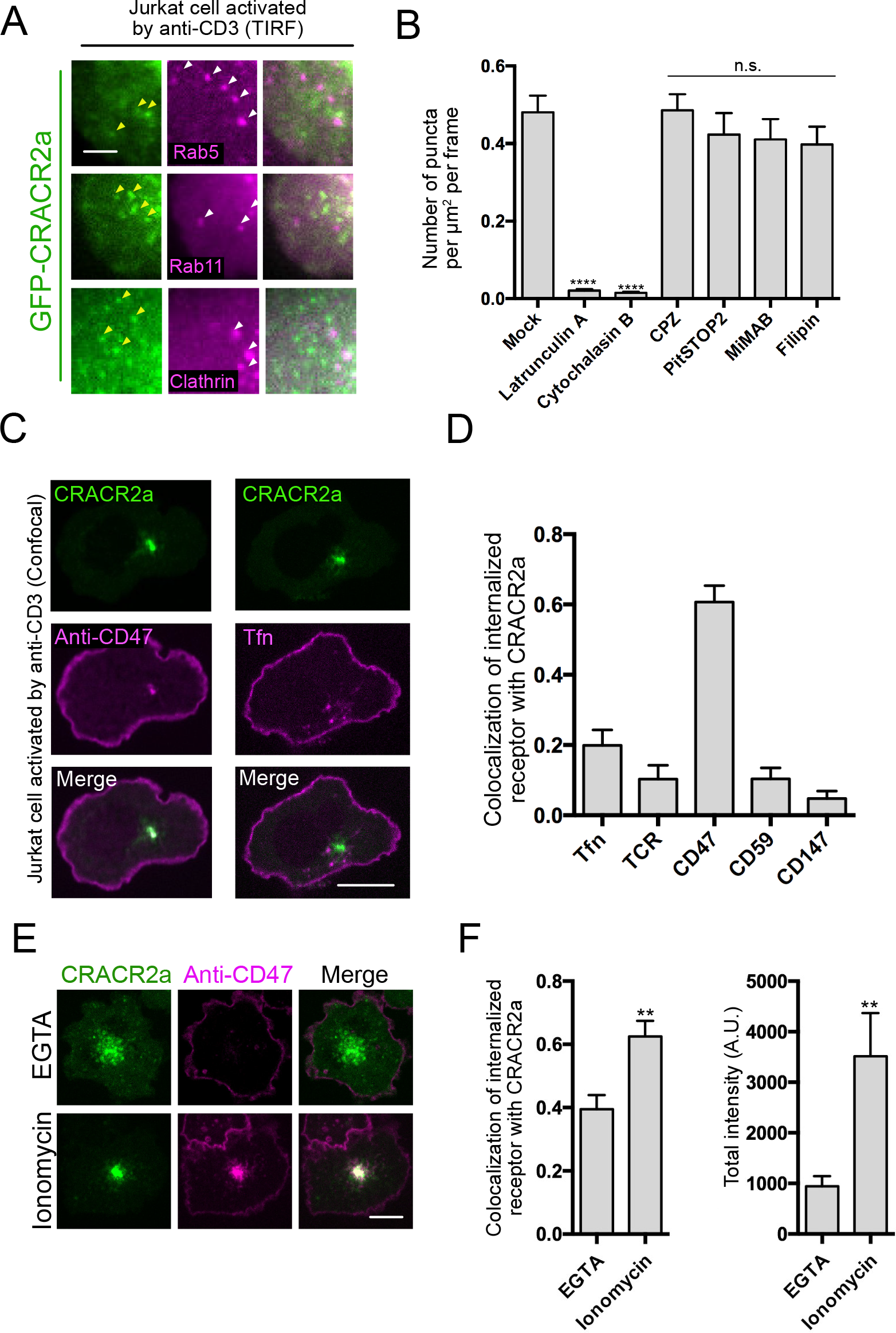
CRACR2a is involved in clathrin independent endocytic trafficking. (**A**) CRACR2a puncta (indicated by yellow arrowheads) do not colocalize with Rab5, Rab11 or clathrin-coated pits (indicated by white arrow heads). Scale bar: 3 *μ*m. (**B**) Quantifications of the number of CRACR2a puncta observed in activated Jurkat cells when treated with different inhibitors. n = 20-30 cells from three independent experiments. Error bar indicates SEM. **** p <0.0001, one way ANOVA with Dunnett’s multiple comparisons test comparing each sample with mock (DMSO) treated sample. (**C**) CD47 but not Transferrin (Tfn) is internalized into CRACR2a compartment at the MTOC. Scale bar: 10 *μ*m. (**D**) Quantification of the fraction of internalized receptor that colocalizes with the CRACR2a compartment. n= 18-25 cells from three independent experiments. (**E**) Ionomycin stimulates CD47 internalization into CRACR2a compartment. (**F**) Left: quantifications of the fraction of internalized CD47 in CRACR2a compartment in EGTA or ionomycin condition; right: quantifications of the total intensity of internalized CD47 in EGTA or ionomycin condition. n = 17-22 cells from three independent experiments. Error bar indicates SEM. ** p <0.01, Student’s T test.

We next sought to identify cell surface proteins that might be internalized into the CRACR2a membrane compartment. We selected a set of candidate molecules, with distinct membrane topologies, that have been shown previously to be internalized through different endocytic pathways (Boucrot et al., 2015; Maldonado-Báez et al., 2013) (Supplementary Table 1). We employed an antibody/ligand internalization assay to examine the endocytosis of cell surface molecules. Briefly, endocytosis was suppressed by incubating cells on ice, and cell surface proteins were stained using fluorescent-conjugated antibodies or with fluorescent transferrin.

Cells were then warmed to 37°C rapidly and stimulated with immobilized anti-CD3 antibodies to promote CRACR2a mediated endocytosis and transport to the MTOC. The amount of internalized antibody (or transferrin) within the CRACR2a compartment was then determined by confocal imaging. Among the candidates tested, only the multi-pass transmembrane protein CD47 was strongly internalized into the CRACR2a compartment (Fig. 5C, 5D). Stimulation of Jurkat cells with ionomycin alone also promoted internalization of CD47 into the CRACR2a compartment at the MTOC (Fig. 5E, 5F). Based on these results, we propose that CRACR2a is involved in a new type of calcium-stimulated, clathrin-independent endocytic pathway, and that CD47 is one of the cargoes internalized through this pathway.

## Discussion

In this study, we provide evidence that the Rab GTPases CRACR2a and Rab45 are activating adaptors for dynein. Several Rab GTPases have been shown to directly bind to dynein adaptors. Rab6, for example, binds to the C-terminal coiled-coil of the adaptor BicD, which then uses its N-terminal coiled-coil to engage dynein-dynactin (Matanis et al., 2002; Urnavicius et al., 2015; Huynh and Vale, 2017). Rab11FIP3 also has N-terminal EF-hands and coiled-coil for dynein activation, while containing a C-terminal Rab11 binding domain (Horgan et al., 2010; McKenney et al., 2014). In these examples, the Rab domains themselves do not appear to contribute directly to binding of dynein-dynactin; instead they mainly act to recruit the adaptors to specific membrane cargoes (Reck-Peterson et al., 2018). Our evidence indicates that CRACR2a and Rab45 represent gene fusions of a coiled-coil dynein adaptor with a Rab GTPase. We also demonstrate that the dynein adaptor function of CRACR2a is regulated by calcium and that calcium release that accompanies T cell activation results in CRACR2a-dynein activation. Finally, we show that CRACR2a-mediated dynein transport also plays a role in extracting clathrin-independent endosomes from the cell cortex and then transporting these vesicles towards the MTOC.

### CRACR2a is a novel calcium-regulated dynein adaptor protein

Rab45 and CRACR2a, together with Rab11FIP3 and the Ninein family, constitute a group of adaptor proteins that contain EF hand motifs and coiled-coil domains. Recent structural studies have demonstrated that the coiled-coil domains play crucial roles in bridging the interaction between dynein and dynactin (Urnavicius et al., 2018, 2015; Zhang et al., 2017). In contrast, the role of the EF hands in these proteins is less clear. Domains outside of the coiled-coil region in adaptor proteins could facilitate or regulate their interactions with dynein-dynactin. The Hook domain of Hook3, for example, directly binds to the dynein light-intermediate chain (Schroeder and Vale, 2016). It is possible that the EF hands in Rab45 and CRACR2a play a similar role. Moreover, EF hands can assume different conformations depending on their calcium binding state (Yap et al., 1999). Interactions between EF hands and their binding partners are therefore often calcium regulated (Lewit-Bentley and Réty, 2000). Our study establishes CRACR2a as the first dynein adaptor protein that responds to calcium changes in vitro and in cells. CRACR2a, therefore, can serve to connect calcium signaling to the regulation of dynein function. We demonstrate that, in T cells, this allows dynein-dependent intracellular transport to be coupled to T cell activation.

### CRACR2a-dynein mediates endocytic transport at the T cell immunological synapse

We observed that CRACR2a localizes to intracellular compartments surrounding the MTOC; interestingly calcium elevation induced by TCR triggering causes a strong compaction of CRACR2a compartments towards the MTOC, which is consistent with an activation of dynein-dynactin motility. Such calcium-induced MTOC clustering was not observed for Rab8A positive trans-Golgi network or Rab11 labeled recycling endosomes, suggesting that CRACR2a localizes to a unique membrane compartment that undergoes calcium-dependent repositioning. We also found one cell surface protein, CD47, that becomes internalized and co-localizes with CRACR2a. In contrast, transferrin receptor, an established cargo for clathrin-mediated endocytosis, does not traffic to CRACR2a compartments. Collectively, these results suggest that the CRACR2a intracellular compartment is derived from a non-clathrin endocytic process and is distinct from Rab8A and Rab11 compartments. The exact nature and function of CRACR2a compartment warrants further investigation.

We observed that CRACR2a forms numerous diffraction-limited puncta at the cell cortex when T cells become activated. These puncta are associated with the actin network and are propelled towards the cell center by the retrograde flow of actin. We further found that CRACR2a puncta colocalize with dynein, and often abruptly disappear from the TIRF field. Confocal imaging suggests that they are pulled into the interior of the cell by dynein-driven transport along microtubules (Fig. 4B). These observations suggest a model in which CRACR2a puncta are first nucleated at the plasma membrane and then become internalized through endocytosis. However, due to the limited Z resolution of TIRF imaging, we cannot determine the precise time when internalization events occur. This form of endocytosis does not co-localize with clathrin and is not affected by pharmacological inhibition of clathrin-coated pit assembly.

Our results also suggest a role of dynein-dynactin and microtubules in some early steps of CRACR2a associated endocytosis (Figure 6), either in mechanical pulling that aids in the formation of the endosomal vesicle itself (Simunovic et al., 2017), or in extracting the vesicle from actin-rich cortex and delivering it to the cell interior. In support of this general idea, CRACR2a puncta are unable to detach from the actin cortex in the absence of microtubule-based transport and instead accumulate underneath the plasma membrane to form a disk-like structure (Fig. 4F, 4G). In contrast to the many studies documenting involvement of actin in endosome formation (Mayor et al., 2014; Ferreira and Boucrot, 2017), a role for microtubules in early events of endocytosis is less well established. An earlier study showed that inhibition of dynein function or disruption of microtubule suppresses a form of chorea toxin-induced tubular endosome formation (Day et al., 2015). This study speculated that a dynein-mediated pulling force generation might promote the membrane invagination required for endocytosis, although whether dynein localizes to these tubular invaginations was not examined. Our data show that dynein and a specific dynein adapter (CRACR2a) are localized at sites of endocytosis in T cells and that microtubule-based transport is required for the movement of nascent vesicles from the actin cortex to the cell interior. More details on the role that dynein plays in this form of endocytosis awaits further studies. However, CRACR2a should provide a useful tool to tease apart this very specific role of dynein.

**Figure 6.**
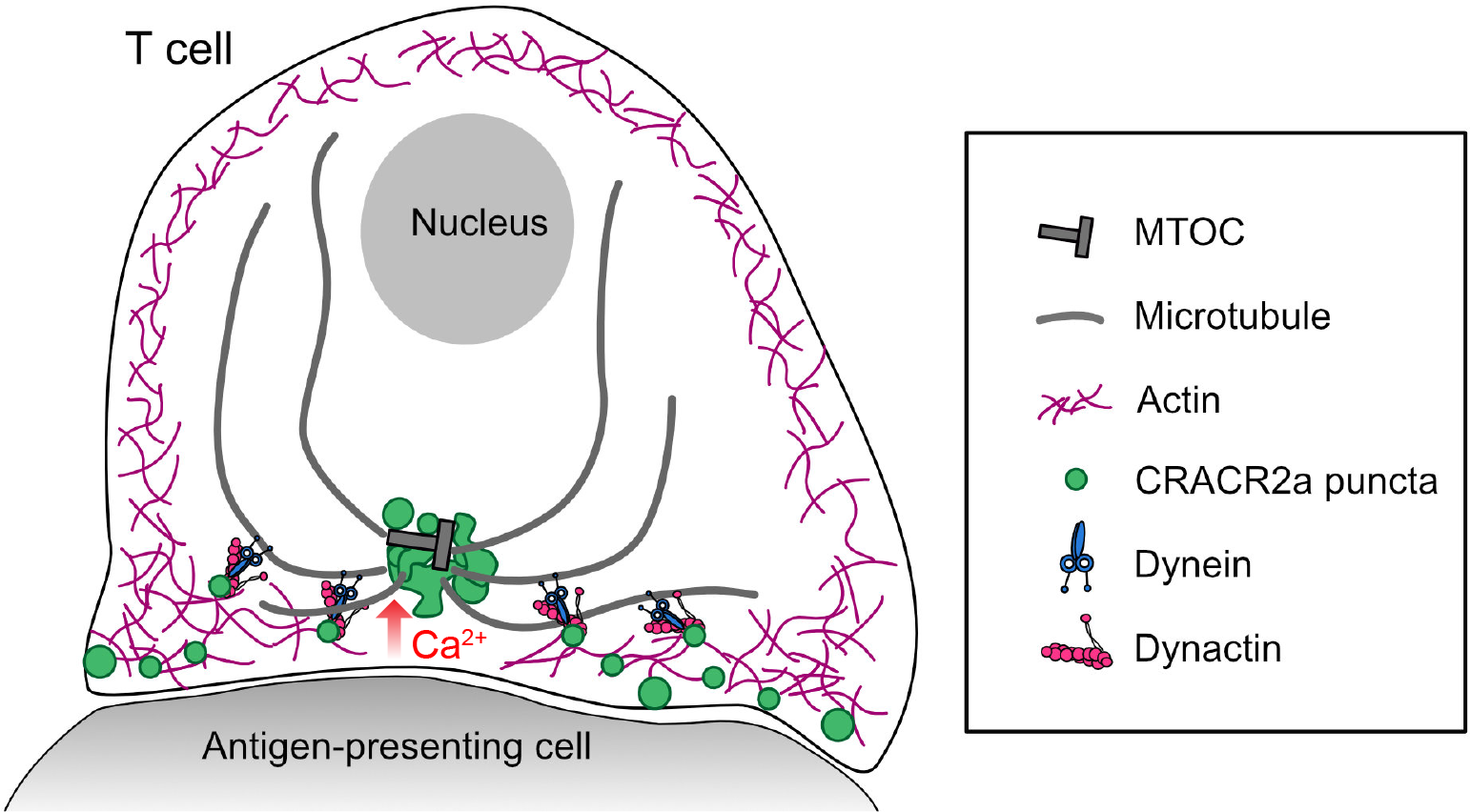
A schematic of CRACR2a-dynein mediated endocytosis during T cell activation. Activation of a T cell by an antigen presenting cell (APC) causes influx of calcium, which promotes the formation of CRACR2a endosomes at the immunological synapse. Activation of dynein-dynactin on CRACR2a endosomes facilitates their detachment from the actin cortex and initiates microtubule minus-end directed transport.

What is the function of CRACR2a in T cells? A previous study showed that siRNA knockdown of CRACR2a blunted TCR signaling, indicating that CRACR2a plays an important role in T cell signal transduction (Srikanth et al., 2016). It was suggested that CRACR2a activates the Rac/Cdc42 guanine nucleotide exchange factor Vav1 through its proline-rich region. While this model is not contradictory with our finding, it is likely that CRACR2a also regulates the endocytosis and/or recycling of T cell signaling molecules. In our initial candidate-based screen of receptor internalization, we found that CD47 is one of the cargoes for CRACR2a mediated endocytosis. CD47 is a ubiquitously expressed transmembrane protein that has diverse functions in regulating cell adhesion, migration and phagocytosis (Brown and Frazier, 2001; Matozaki et al., 2009). The role of CD47 in T cells remains poorly studied, though earlier reports suggested that CD47 plays regulatory roles in integrin-mediated adhesion and T cell migration (Azcutia et al., 2013; Piccio et al., 2005; Azcutia et al., 2017). Future investigation of the endocytosis process mediated by CRACR2a will help to elucidate its role in T cell function.

## Acknowledgments

We thank N. Stuurman for help with microscopy and image analysis, and A. Weiss and W. Lu for helpful discussion and for providing T cell related reagents. We would also like to thank the Vale lab members for discussions and critiques of the manuscript. This work was funded by the National Institutes of Health (5R35GM118106-02, R.D.V.) and the Howard Hughes Medical Institute.

## Author Contributions

Y. W. and R.D.V. conceived of the study and designed the experiments. W. H. and T. S. performed the peroxisome assays and initial biochemical purifications and analyzed the data. Y. W. performed all the other experiments and analyzed the data. Y. W. and R.D.V. wrote the manuscript and all co-authors provided feedback.

## Video Legends

**Video 1. Confocal time-lapse imaging of GFP-CRACR2a intracellular compartments in Jurkat T cells** Jurkat cells stably expressing GFP-CRACR2a were allowed to settle on glass without coating. Video was captured by spinning disk confocal microscopy with 4 frames per sec acquisition rate. Scale bar: 5 *μ*m.

**Video 2. Dynamics of CRACR2a puncta at the T cell immunological synapse** Jurkat cells stably expressing GFP-CRACR2a were allowed to settle on anti-CD3 coated glass. Video was captured by TIRF microscopy at 1 frame every 4.9 sec. Scale bar: 5 *μ*m. On the right is a 2x magnified view of a section of the cell. Scale bar: 5 *μ*m.

**Video 3. CRACR2a puncta migrate with actin retrograde flow** Jurkat cells stably expressing GFP-CRACR2a (Green) and BFP-LifeAct (Magenta) were allowed to settle on anti-CD3 coated glass. Video was captured by TIRF microscopy at 1 frame every 3.5 sec. Scale bar: 5 *μ*m.

**Video 4. Latrunculin treatment disrupts the retrograde movement of CRACR2a puncta** Jurkat cells stably expressing GFP-CRACR2a were allowed to settle on anti-CD3 coated glass. Latrunculin A (LatA) was added at 200 sec timepoint at 2 *μ*M final concentration. Video was captured by TIRF microscopy at 1 frame per 2 sec. Scale bar: 5 *μ*m.

**Video 5. Nocodazole treatment causes accumulation of CRACR2a puncta at central region of the immunological synapse.** Jurkat cells stably expressing GFP-CRACR2a were treated with 5 *μ*M nocodazole for 20 min and then allowed to settle on anti-CD3 coated glass. Video was captured by TIRF microscopy at 1 frame per 4.8 sec. Scale bar: 5 *μ*m.

## Supplementary Figure Legends

**Figure S1 (related to Figure 1).**
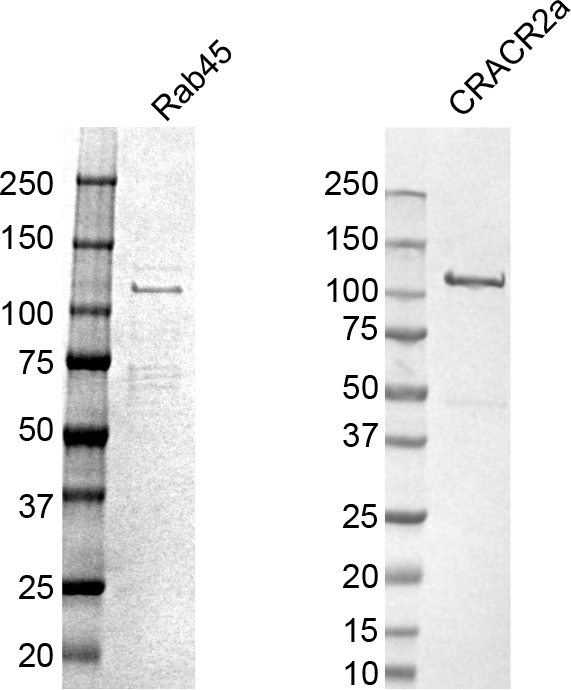
SDS PAGE of purified GFP tagged Rab45 and CRACR2a. The molecular weight of the protein standard loaded is labeled on the left of each panel.

**Figure S2 (related to Figure 2).**
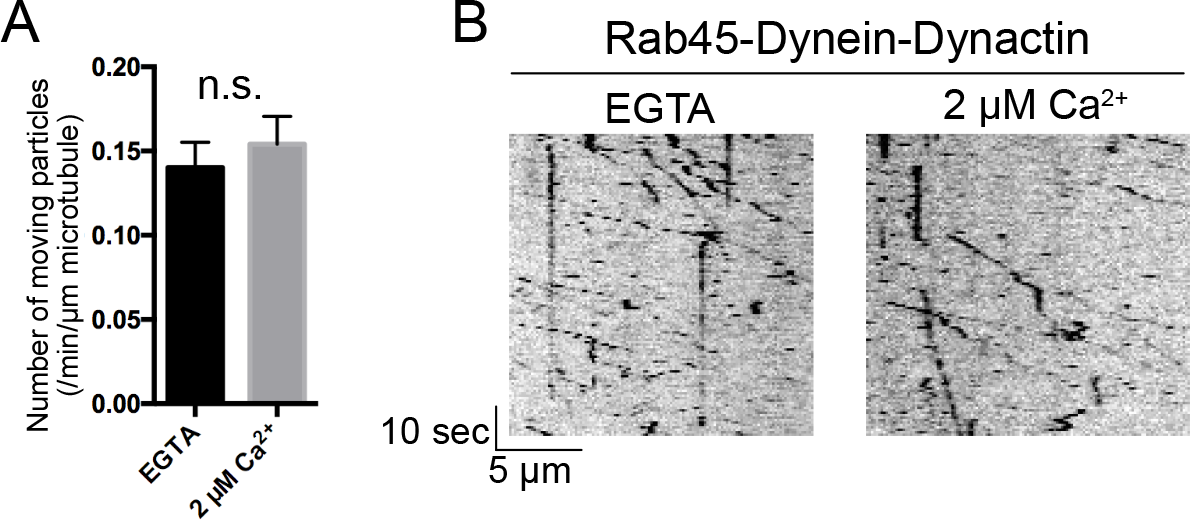
The dynein adaptor function of Rab45 is not regulated by calcium. (**A**) Quantification of Rab45-dynein-dynactin motility in calcium-depleted buffer (EGTA) or buffer with 2 *μ*M free calcium. Data are from three replicates, each measuring at least 20 microtubules. Error bar: SEM. N.S. indicates not statistically significant, student’s T-test. (**B**) Sample kymographs of Rab45-dynein-dynactin moving on microtubules in EGTA or 2 *μ*M calcium condition.

**Figure S3 (related to Figure 3).**
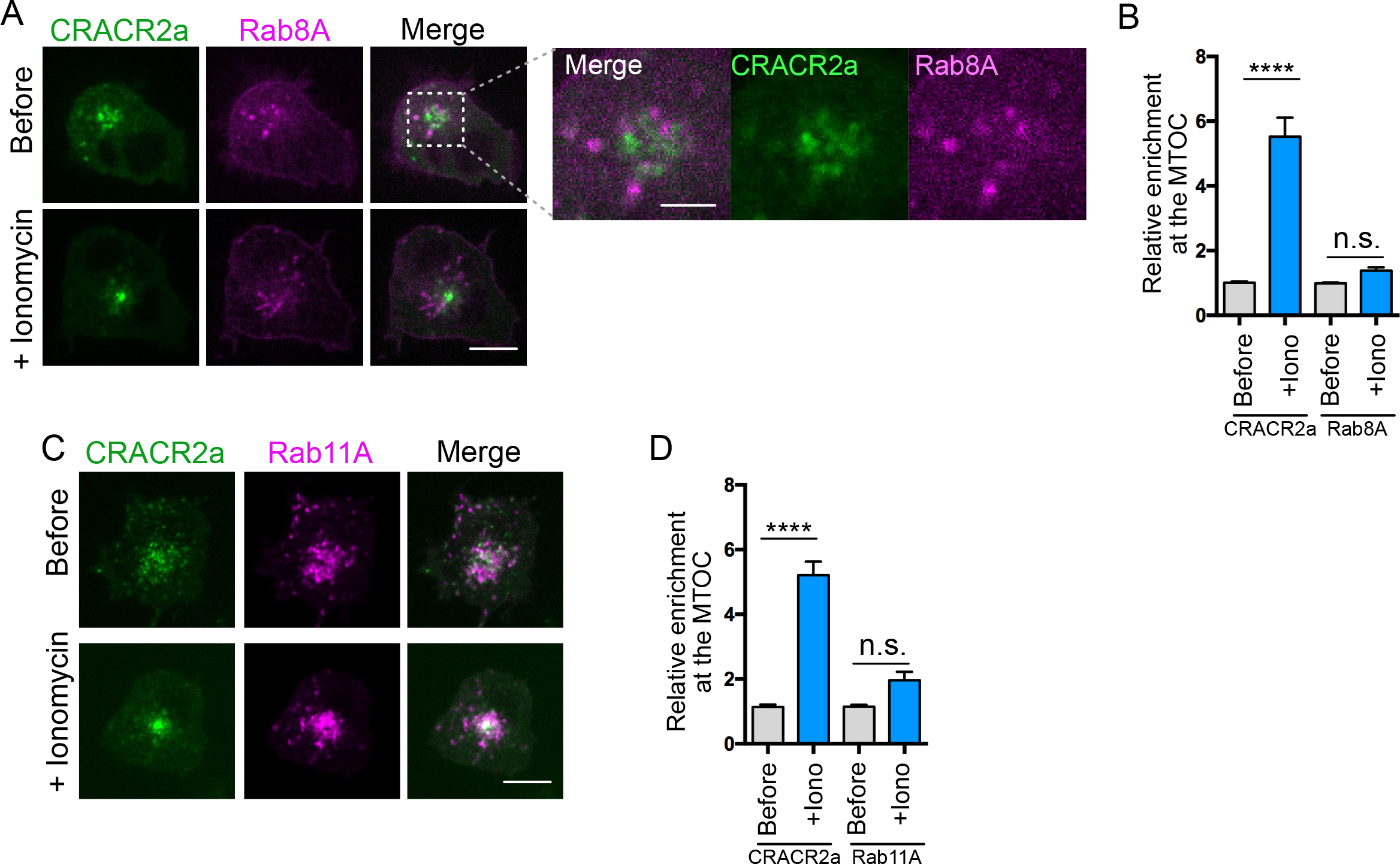
CRACR2a does not colocalize with Rab8 or Rab11 in T cells. (**A**) Subcellular localization of GFP-CRACR2a and Rab8A in T cells before or after stimulation of the cell with Ionomycin/Ca^2+^. Scale bar: 5 *μ*m. A cropped region shows Rab8A and CRACR2a do not overlap (scale bar: 2 *μ*m). (**B**) Quantifications of the levels of relative enrichment at MTOC for CRACR2a or Rab8A. n = 18-22 cells from three independent experiments. Error bar indicates SEM. **** p<0.0001, one way ANOVA analysis with Sidak’s multiple comparisons test. (**C**) Subcellular localization of GFP-CRACR2a and Rab11A in T cells before or after stimulation of the cell with Ionomycin/Ca^2+^. Scale bar: 5 *μ*m. (**D**) Quantifications of the levels of relative enrichment at MTOC for CRACR2a or Rab11A. n = 16-23 cells from three independent experiments. Error bar indicates SEM. **** p<0.0001, one way ANOVA analysis with Sidak’s multiple comparisons test.

## Supplementary Table Legends

**Supplementary Table 1.** Candidate cell surface proteins tested for internalization into CRACR2a compartment.

|  | **Transferrin Receptor** | **TCR** | **CD47** | **CD59** | **CD147** |
| --- | --- | --- | --- | --- | --- |
| Type | Type II single-transmembrane protein (dimer) | Multi-chain receptor complex | Five-pass transmembrane protein | GPI-anchored cell surface protein | Type II single-transmembrane protein |
| Endocytic pathway | CME | CME and CIE | CIE?* | CIE | CIE |

\* listed as CIE cargo in Boucrot et al., 2015 but no primary data could be found.
CME: Clathrin-mediated endocytosis
CIE: Clathrin-independent endocytosis

## Materials and Methods

### DNA constructs, antibodies and chemical inhibitors

The cDNA for human Rab45 (NM_152573.3) and CRACR2a (NM_001144958.1) were synthesized by Genscript. Detailed information about the various DNA constructs used can be found in Supplementary Excel file, “Constructs” tab. Information about antibodies and chemical inhibitors used in the study can be found in “Antibodies” tab and “Inhibitors” tab, respectively.

### Protein purification

Rab45 and CRACR2a expressed in bacteria are prone to degradation. To enrich for full-length proteins, we adopted a tandem affinity purification strategy. Briefly, Rab45 and CRACR2a constructs in pET28 vectors were transformed into the Escherichia coli strain BL21 RIPL (Agilent). Bacterial culture was grown in Terrific Broth at 37°C until growth reached ~2.0 OD_600_. The temperature was then lowered to 18°C and the culture was induced overnight with 0.5 mM Isopropyl β-D-1-thiogalactopyranoside (IPTG). Bacterial pellets were resuspended in Ni-A buffer (50 mM Tris-HCl pH 8.0, 500 mM NaCl, 2 mM MgCl_2_, 5% Glycerol, 10 mM Imidazole and 3 mM 2-Mercaptoethanol). Cells were lysed using an Emulsiflex press (Avestin) and clarified at 18k RPM using SORVALL SS-34 rotor for 60 min at 4°C. Lysates were filtered through a 0.45 *μ*m filter before loading to a Histrap FF column (GE Healthcare). Column was then washed with 20 column volumes (CV) of Ni-A buffer. Bound protein was eluted with 10 CV Ni-B buffer (20 mM Tris-HCl pH 8.0, 300 mM NaCl, 5% Glycerol, 500 mM Imidazole and 3 mM 2-Mercaptoethanol). Eluate was then directly loaded to a Streptrap HP column (GE Healthcare). Column was washed with 20 CV wash buffer (20 mM Tris-HCl pH 8.0, 300 mM NaCl, 2 mM MgCl_2_, 5% Glycerol, 2 mM DTT) and the bound protein was eluted with 10 CV elution buffer (20 mM Tris-HCl pH 8.0, 300 mM NaCl, 2 mM MgCl_2_, 5% Glycerol, 3 mM Desthiobiotin, 2 mM DTT). Eluted proteins were further purified with a Superose 6 10/300GL (GE healthcare) gel filtration column.

Peak fractions were pooled and concentrated and then flash frozen in liquid nitrogen. Dynein and dynactin were prepared from RPE-1 cells, as previous described (Huynh and Vale, 2017).

### EGTA buffered calcium solution

In order to maintain free calcium concentration at physiologically relevant level (100 nM ~ 10 *μ*M) it is essential to use a EGTA:Ca^2+^ buffer system, which is prepared by mixing EGTA and CaCl2 at a calculated ratio to achieve desired free calcium concentration (Bers et al., 2010). We followed the protocol described by Bers et al., 2010 and used the Maxchelator tool that the authors developed (http://maxchelator.stanford.edu/index.html) to calculate the ratio of EGTA and CaCl_2_ needed to maintain the desired calcium concentration.

### Dynein-dynactin pull down assay

Pull-down of dynein/dynactin was performed as previously described (Huynh and Vale, 2017). Briefly, dynein/dynactin isolated from RPE1 cells were mixed with 20 nM purified Rab45 or CRACR2a and 15 μL pre-equilibrated Streptactin Sepharose beads (GE Healthcare) in 300 μL binding buffer (50 mM HEPES pH 7.4, 20 mM NaCl, 1 mM Mg-Acetate, 10% Glycerol, 0.1% NP40, 2 mM DTT). The mixture was incubated at 4°C for 1 hr. Beads were pelleted at 1000x rcf for 2 min and washed 4 times with 500 μL binding buffer. The proteins were eluted by boiling in SDS loading buffer for 5 min, and loaded onto SDS PAGE gel. The amount of CRACR2a and Rab45 in the pull-down samples were visualized by Coomassie staining. The amount of dynein and dynactin were detected via western blot with antibodies against dynein heavy chain (DHC) and the p150 subunit of dynactin, respectively. In the pull-down assay shown in Fig. 2A, 2 mM EGTA or EGTA:Ca^2+^ was added to the binding buffer to deplete calcium or to maintain a free calcium concentration of 2 *μ*M. Information about antibodies used can be found in Supplementary Excel file.

### Single molecule motility assay

Microtubules were prepared as previously described (Huynh and Vale, 2017). Briefly, unlabeled tubulin, biotinylated tubulin and Alexa-640 labeled tubulin were mixed at a ratio of ~5:1:1 in BRB80 (80 mM PIPES, pH 6.8, 1 mM EGTA, 1 mM MgCl2). GTP was added to 5 mM final concentration and the mixture was incubated at 37°C for 10 min. Taxol was then added to 20 μM and microtubules were allowed to polymerize overnight. Before use microtubules were spun over a 25% sucrose cushion in BRB80 with 10 μM taxol at 20,000 g for 10 min, and resuspended in BRB80 with 10 *μ*M taxol but without EGTA.

For In vitro motility assays, purified Rab45 or CRACR2a were mixed with dynein-dynactin at a 5:1 ratio in a 50 *μ*L reaction volume in assay buffer (50 mM HEPES pH 7.4, 2 mM magnesium acetate, 10% glycerol, 2 mM DTT) along with 0.1 mg/ml Biotin-BSA, 0.5% pluronic acid F-127, and 0.2 mg/ml κ-casein. In the motility assay shown in Fig. 2B, EGTA:Ca^2+^ buffer was added to the assay buffer to maintain the desired free calcium concentration. The mixture was incubated on ice for 1 hr. Flow chambers with attached microtubules were prepared as described (Schroeder and Vale, 2016). A 1:5 dilution of the dynein-dynactin-adaptor complex was then added in the presence of 1 mM ATP and the Trolox/PCD oxygen scavenging system (Aitken et al., 2008). Total internal reflection fluorescence imaging of single molecule was performed on a Nikon Eclipse TE200-E microscope equipped with an Andor iXon EM CCD camera, a 100x 1.49 NA objective, and MicroManager software (Edelstein et al., 2014). Exposures conditions were 200 ms per frame with 1 sec interval for 150 frames. Kymographs were created for randomly selected microtubules using Fiji (Schindelin et al., 2012), and the number of processive particles were counted manually.

### Cell culture and transduction

U2OS cells and HEK293T cells were cultured in DMEM medium containing 10% FBS and penicillin/streptomycin/L-glutamine. Jurkat cells were cultured in RPMI1640 medium containing 10% FBS and penicillin/streptomycin/L-glutamine. Lentiviral transduction was used to create cell lines stably expressing GFP-CRACR2a, and GFP positive cells were selected by FACS. The plasmids expressing mCherry-Rab5A, mCherry-Rab11A, mCherry-Clathrin light chain and BFP-Rab8A were transiently transfected using TransIT-Jurkat (Mirus Bio) following the manufacture’s protocol.

### Peroxisome assay

Unlike other dynein adaptors, Rab45 and CRACR2a contain a Rab GTPase domain that associates with intracellular membranes through its C-terminal geranylgeranylation. To prevent concurrent localization of Rab45 and CRACR2a to peroxisomes and non-peroxisomal membrane compartments, which would complicate the analysis, we deleted their C-terminal CCX prenylation motifs and cloned it into the pmCherry-FKBP plasmid. The assay was performed as previously described (Schindelin et al., 2012). Briefly, U2OS cells were co-transfected with mCherry-adaptor-FKBP and PEX-GFP-FRB plasmids. 24 hr post transfection, cells were treated with 0.25 *μ*M rapamycin for 50 min, and for the last 10 min, cells were stained with CellMask DeepRed. After washing with PBS twice, cells were fixed with 4% paraformaldehyde for 15 min at room temperature. Images were acquired on a Nikon Eclipse TE200-E spinning disk confocal microscope equipped with an Andor iXon EM CCD camera, a 40x 0.95 NA objective, and MicroManager software (Edelstein et al., 2014). We manually quantified the number of cells exhibiting a clustered peroxisome phenotype, which is defined as all of the peroxisome signal in the GFP channel coalescing into a cluster at the cell center.

### T cell activation and calcium stimulation

Glass-bottom 96-well plates (MatriCal) were coated with anti-CD3ε antibody (clone OKT3) or with IgG isotype control at room temperature for 2-4 hr or at 4°C overnight. In Fig. 5D to monitor TCR internalization, cells were activated by a different anti-CD3 antibody (clone UCHT1), which does not compete with the Alexa 647 conjugated antibody used for labeling TCR. Wells were washed once with PBS. Jurkat T cells were resuspended in RPMI-1640 media (no Phenol-red, with 25 mM HEPES pH 7.4) and rested for 30 min before being dropped onto antibody-coated wells. For ionomycin stimulation, Jurkat cells were resuspended in Hank’s Balanced Buffer Solution with 25 mM HEPES pH 7.4 and added to wells without coating. Cells were allowed to settle for 15 min, and then ionomycin and CaCl_2_ were added to final concentrations of 0.5 *μ*M and 1 mM, respectively.

### Confocal imaging of CRACR2a endosomes and perinuclear compartments

Jurkat cells expressing GFP-CRACR2a were stimulated as described above. For SiR-tubulin staining, cells were stained with 200 nM SiR-tubulin in their growth medium for 4 hr at 37°C. Cells were imaged on a Nikon Eclipse TE200-E spinning disk confocal microscope equipped with an Andor iXon EM CCD camera, a 100x 1.49 NA oil objective, and MicroManager software (Edelstein et al., 2014). To calculate the relative enrichment of the CRACR2a compartment at the MTOC, a series of z sections at 0.3 *μ*m z interval was taken. The location of the MTOC was identified through the SiR-tubulin channel. The relative enrichment at the MTOC was quantified using Fiji by measuring the total intensity of GFP signal within 1 *μ*m radius of the MTOC, which was then normalized by the total intensity of GFP in the 1 – 5 *μ*m radius around the MTOC. In Fig. 4D cells expressing BFP-LifeAct and GFP-CRACR2a were labeled with 200 nM SiR-Tubulin for 4 hrs. A single frame of LifeAct and SiR-Tubulin was acquired and the cells were then imaged continuously using a 488 nm laser at 6.7 frames per second acquisition rate.

### TIRF imaging of CRACR2a endosome at the immunological synapse

Jurkat cells expressing GFP-CRACR2a were stimulated as described above. TIRF imaging was performed on a Nikon Eclipse TE200-E microscope equipped with a motorized TIRF arm, a Hamamatsu Flash 4 camera, a 100x 1.49 NA oil objective, and MicroManager software (Edelstein et al., 2014). We found that the GFP-CRACR2a puncta have a very low signal to noise ratio, possibly due to a high level of diffusive GFP signal present at the cell membrane. We used relatively low laser power and long exposure time (800ms) with 2-4 second imaging interval to improve the signal-noise ratio and to reduce photobleaching. To quantify the number of CRACR2a puncta formed, we used the ImageJ plug-in “Spot Counter” (http://imagej.net/SpotCounter) with “BoxSize” set to 3 and “Noise tolerance” set to 100. The number of detected spots was quantified over 20 frames and the average number or spots per frame per micron was calculated. One or two ROIs at the periphery of the cell were selected for quantification. For data shown in Fig. 5B, various chemical inhibitors were added and cells were treated for 5 min before being imaged using TIRF-M. For data shown in Fig. 4F and 4G, Jurkat cells were treated with 5 *μ*M nocodazole for 30 min in RPMI1640 before being added to antibody-coated wells. Detailed information regarding inhibitors and concentrations used can be found in Supplementary Excel file.

### Antibody internalization assay

For data shown in Fig. 5C, Jurkat cells were washed once with RPMI1640 (no Phenol-red, with 25 mM HEPES pH 7.4) to remove serum. Cells were then incubated on ice for 10 min to inhibit endocytosis, before stained with Alexa Fluor 647 labeled antibodies or transferrin for 20 min on ice. Cells were then washed twice with cold RPMI1640 before being added to anti-CD3 coated wells with pre-warmed RPMI1640. 25-30 min after addition of the cells to the wells, z-sections were acquired using a spinning disk confocal microscope at 0.3 *μ*m z interval. For data shown in Fig. 5E, cells were stained with Alexa Fluor 647 conjugated anti-CD47 as described above, except 1 mM EGTA was added to deplete calcium from the medium. Cells were allowed to settle on non-coated glass for 20 min and confocal z-sections were acquired. Cells were then treated with 0.5 *μ*M ionomycin and 2 mM CaCl_2_ for 5 min and confocal z-sections were taken. The JACoP plug-in in Fiji was used for quantifications of the colocalization of internalized receptor in CRACR2a compartments (Bolte and Cordelières, 2006). A representative z section was selected. After background subtraction using autothresholding, the colocalization was calculated using the M1 & M2 coefficients analysis. We defined the GFP-CRACR2a channel as channel A, and Alexa Fluor 647 channel as channel B. The M2 coefficient was calculated as follows (Bolte and Cordelières, 2006):

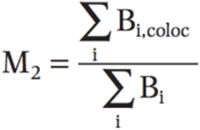

B_i_ is the intensity of pixel i in channel B; A_i_ is the intensity of pixel i in channel A

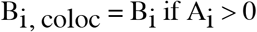

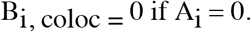

In this experiment the M2 coefficient represents the amount of internalized receptor within the CRACR2a compartment normalized by the total amount of internalized receptor.

